# Repulsion from slow-diffusing nutrients improves chemotaxis towards moving sources

**DOI:** 10.1101/2024.01.25.577248

**Authors:** Blox Bloxham, Hyunseok Lee, Jeff Gore

## Abstract

Chemotaxis, or the following of chemical concentration gradients, is essential for microbes to locate nutrients. However, microbes often display paradoxical behaviors, such as *Escherichia coli* being repelled by several amino acids. Here, we explore chemotaxis towards a moving source and demonstrate that when multiple nutrients are released from the source repulsion from certain nutrients actually improves chemotaxis towards the source. Because a moving source leaves most of the nutrient plume behind it, simply following the concentration gradient results in aiming behind the source and failing to intercept it. However, when attraction to a fast-diffusing nutrient and repulsion from a slow-diffusing nutrient are combined, motion in a new direction emerges and the chance of intercepting the source is increased up to six-fold. We demonstrate that this “differential strategy” is robust against numerous variations, including order-of-magnitude increases in the repellent release rate. Finally, we leverage existing data to show that *E. coli* is attracted to fast-diffusing amino acids and repelled by slow-diffusing ones, suggesting it may utilize a differential strategy and providing an explanation for its repulsion from these amino acids. Our results thus illuminate new possibilities in how microbes can integrate signals from multiple gradients to accomplish the difficult chemotactic tasks.

## Introduction

The ability to gather nutrients is essential for any organism to grow. In a spatially heterogeneous world, the acquisition of nutrients can be greatly improved by moving towards locations with the most favorable conditions^1–4^. For microbes, this generally involves following chemical concentration gradients towards increasing concentrations of attractants and away from repellants, a behavior known as chemotaxis^3,5–12^. Attractants can be nutrients diffusing from their source^13–15^ or other indicators of favorable conditions^7^. Repellants are often, but not always^16^, toxins or other harmful compounds or conditions^16–18^. Chemotaxis is essentially a gradient ascent: to find the highest nutrient concentration, microbes simply move in whichever direction it’s locally increasing.

Microbes do, however, display surprising patterns as to which compounds are attractants and which are repellants. For example, *Escherichia coli* is repelled by certain amino acids^16,18^ including some it can utilize as food^18^, as well as other nutrients such as glycerol^19^ and even combinations of attractants^20^. A recent investigation demonstrated that many of the amino acids *E. coli* is repelled by are less effective nutrients for it and in some cases inhibit growth^18^. Anomalies do remain, however, such as tryptophan being one of the amino acids that supports the most growth as the sole nitrogen or nitrogen and carbon source but also being the strongest repellant^18^. Other surprising chemotactic responses include *Pseudomonas aeruginosa* being attracted to lethal antibiotics such as ciprofloxacin and streptomycin, possibly as a strategy to target and invade competing colonies^21^. Another study observed that marine *Alteromonas* and *Pseudoalteromonas* isolates’ chemotactic preferences for the polysaccharide laminarin varied heavily based on the molecular weight of the polymer with some strains displaying their strongest chemotaxis towards high-molecular weight polymers and other strains displaying their strongest chemotaxis towards low-weight polymers and the monosaccharide components^22^. Microbes have also been observed to be sometimes attracted to and sometimes repelled by the same compounds, depending on concentration and other factors^23^. *E. coli’s* repulsion from nutrients and these other intriguing results suggest that microbes may be integrating external signals in sophisticated ways to carefully tune their trajectories.

Natural sources of chemoeffectors (i.e. attractants and repellents) will generally be sources of more than one, and the resulting presence of multiple concentration gradients creates additional opportunities for sophisticated signal processing. When faced with two opposing attractant gradients microbes must choose between the two, with outcomes potentially dependent on previous environmental exposure and resulting receptor expression^24–26^. In the presence of parallel attractant and repellant gradients, microbes must make a similar choice between net attraction or repulsion^27–32^, with chemotaxis to an intermediate distance from the source or a ‘bet-hedging’ split into attracted and repelled subpopulations also being possible outcomes^28^. In some cases, a microbe will only respond to certain chemoeffectors if another is present^20^. On the broader theme of chemotaxis in realistically complex environments, the combination of faster growth on a colony’s edge and chemotaxis towards compounds secreted during growth can significantly boost range expansion^33,34^. However, while recent research has illuminated the importance of carefully considering spatial structure when exploring chemotaxis, previous reports appear to have only considered chemoeffector gradients that are roughly parallel or antiparallel^24,26,28,29,31^, leaving the effect of combining nonparallel gradients largely unexplored. More broadly, how the surprising chemotactic behaviors of microbes and the integration of multiple signals come together to solve complex, real-world navigation tasks presents exciting opportunities for exploration.

In addition to being spatially structured, the world is in constant motion, so chemotaxis will often involve intercepting a moving source. An important example is marine microbes intercepting sinking particulate organic matter, where rates of successful intercepts have important implications for the global carbon cycle^35^. These particles may sink faster than the microbe can swim^36–38^, so small differences in trajectory can make the difference between a successful intercept and getting left behind unable to ever catch up^39^. Chemotaxis towards a moving source could also take the form of one microbe attempting to intercept another for the purposes of predation or cross feeding^40–42^. Even for a stationary source, the presence of a fluid flow leads to an equivalent scenario under a change in reference frame^43,44^. Thus, it is essential to consider the performance of chemotactic strategies in the case of a moving source to fully understand nutrient acquisition in natural environments, and doing so may yield insight into the observed complexities of chemotactic behavior.

Here, we demonstrate how repulsion from a nutrient can actually improve chemotaxis towards a moving source. Specifically, we show that combining attraction to a fast-diffusing nutrient and repulsion from a slow-diffusing nutrient through heterogeneous receptor clusters obeying Monod-Wyman-Changeux dynamics significantly increases the likelihood of intercepting a moving source. Indeed, for small sources moving about twice as fast as the microbe can swim, the range of initial positions from which the microbe can successfully intercept the source is more than six-fold larger under this differential strategy than under purely attractive chemotaxis. We further show that the differential strategy is robust to numerous variations in microbe and source characteristics, including order-of-magnitude increases in the rate at which repellent is released from the moving source. We conclude by highlighting that *E. coli* is attracted to fast-diffusing amino acids and repelled by slow-diffusing ones – exactly as needed to implement this strategy. Our results thus present a novel mechanism for microbes to accomplish a difficult task and suggest an explanation for surprising experimental observations.

## Results

### Following the gradient of an attractant is a poor strategy for intercepting a moving source

We began by benchmarking the difficulty of intercepting a moving source via purely attractive chemotaxis. We considered a moving source with constant downwards velocity between 0 and 7.5 μm/s releasing an attractant with diffusion coefficient 1000 μm/s^2^ at a constant rate of 1 fmol/s and a microbe that swam at 6 μm/s (an approximate net drift velocity for *E. coli*^45,46^). As the source is in constant motion, it leaves a plume of nutrient behind it as the molecules diffuse from past locations (Fig. 1A). The optimal trajectory to intercept a moving target is to aim in front of it (where it will be at the time of interception; Fig. 1B), but the nutrient concentration lagging behind the source means the concentration gradient always points behind the source (Fig. 1C). This discrepancy only becomes larger as the source velocity increases. For example, when the source is moving 20% faster than the microbe can swim, the direction of the concentration gradient differs from the optimal trajectory by more than 60° on average (Fig. 1C) and by more than 90° at some locations (Fig. 1B). When the source is moving five times faster than the microbe can swim, the average discrepancy increases to more than 90°. Thus, the time-lagged nature of signals from nutrients diffusing from a moving source appears to make intercepting moving sources via chemotaxis an inherently difficult task.

**Fig. 1.**
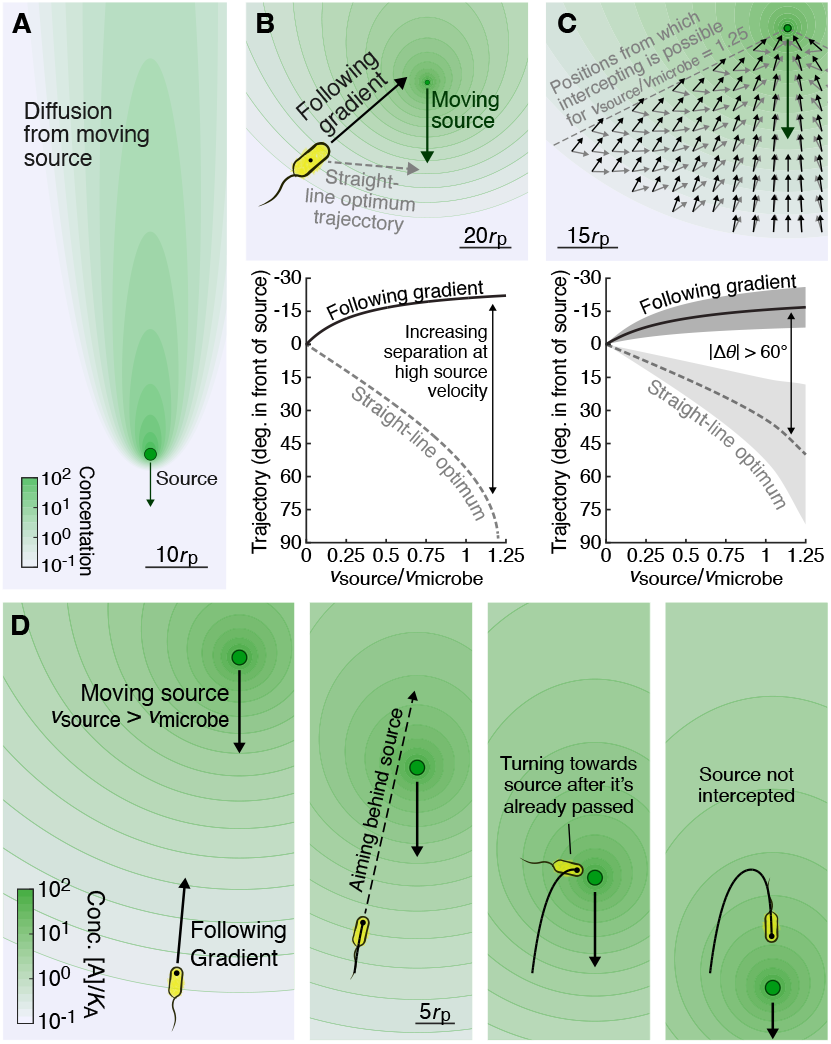
Following the gradient of an attractant leads to inefficient and often unsuccessful interception of a moving source. **A** Cross-section of the plume created by a nutrient diffusing in 3D space at 1000 μm^2^/s from a spherical source particle moving at a constant downwards velocity. Concentration shown in arbitrary units. Scale bar indicates 10 particle radii. The plume is symmetric under rotations around the axis the source is traveling along (Methods). **B** Top: Direction a microbe (not drawn to scale in this nor any other figures) located at the position of the black dot following the concentration gradient would travel in (solid arrow), compared to the optimal straight-line trajectory for the fastest possible intercept (dashed arrow, Methods). Source is a spherical particle of radius 10 μm sinking at 4.5 μm/s and is drawn to scale. Same colormap as **A**. Bottom: How those two directions (defined as angles relative to the straight line to the source’s current position, Methods) change with the source velocity. **C** Top: Same as **B** but now for an array of points. Bottom: Mean and standard deviation of the two directions of travel across the spatial region indicated above (Methods). **D** Example trajectory (in the relevant 2D cross-section in 3D space, Methods) illustrating how the inefficiency of simply following the concentration gradient can prevent a microbe from intercepting a moving source. Because the nutrient concentration is rotationally symmetric about the axis the source is traveling along all motion occurs in the plane containing that axis and the microbe’s initial position (Methods). In this and all later Main Text illustrations, the source is a particle of radius 10 μm moving at 9 μm/s, and the microbe has a maximum speed of 6 μm/s (based on observed net drift velocities for *E. coil*, Methods). Frames show the arrangement of the source and microbe at increments of 15.5 seconds. The nutrient is released from the source at 1 fmol/s, and its concentration is shown as a colormap normalized to the microbe’s receptor binding affinity to the nutrient (100 nM).

To confirm that this geometric challenge can indeed prevent a microbe from successfully intercepting a moving source, we simulated the trajectories of microbes following the gradient of the attractant, which first necessitated assembling a model of microbial chemotaxis. Due to how incredibly well *E. coli*’s chemotactic response has been characterized^7,9,11,26,46–48^ and that many of its key features appear to be conserved across diverse bacteria^49–52^, we chose to use a model and parameters relevant to *E. coli* in the Main Text (while also confirming applicability to marine microbes in Supplementary Figure S3). We used a deterministic approximation in which the microbe swam exactly in the direction of the concentration gradient (but reproduced our key results with stochastic “run-and-tumble” motion in Supplementary Figure S2). The microbe’s velocity depended on the magnitude of the concentration gradient as derived from a Monod-Wyman-Changeux model of chemoeffector binding^10,26,46,53–55^ with a convergence to a maximum velocity of 6 μm/s additionally imposed (Methods). We focused on sources moving faster than the microbe’s maximum velocity to study cases in which a difference in strategy could make the difference between whether the microbe could ever intercept the source at all. The source was a spherical particle whose radius was initially set at 10 μm and velocity at 9 μm/s (consistent with a density of approximately 1.05 g/cm^3^). The attractant was released at a constant rate of 1 fmol/s (in approximate agreement with Kiørboe and Jackson 2001^56^) with the microbe’s receptor binding affinity being *K*_*A*_ = 100 nM^18^. Due to the low Péclet number (0.09), we used a point source approximation for calculating the attractant concentration (Methods) and neglected the effects of fluid flows around the source particle (but reproduce key results with the effect of fluid flows on both concentration plumes and microbe trajectories accounted for in Supplementary Figure S1). Trajectories were calculated in 3D but, due to an azimuthal symmetry of the nutrient plume (Methods), were confined to a 2D plane containing the source’s line of travel and the microbe’s initial position. We thus had a model of microbial chemotaxis to a sinking particle of sufficient simplicity to be tractable and interpretable while still accurately representing microbial chemotaxis and a realistic source.

As an initial demonstration, we used our model to simulate a microbe starting 400 μm below the source’s initial position and 80 μm from the axis the source is traveling along (Fig. 1D). This initial geometry should not, naïvely, pose any particular challenge to the microbe’s attempt to intercept the source: the microbe could, for example, spend just 12 seconds swimming horizontally until it is under the path of the outer edge of particle and then wait another 32 seconds for the particle to arrive. However, the microbe’s trajectory begins with it swimming almost directly upwards, aiming behind the source (Fig. 1D). The microbe does eventually turn downwards but does not do so until after the source has passed, at which point it is too late to ever intercept the source because it is now moving away faster than the microbe can swim (Fig. 1D). Thus, a microbe following this simple “purely attractive” chemotactic strategy is unable to intercept a moving source that should seemingly not be at all challenging to intercept.

### Adding repulsion from a slow-diffusing nutrient allows microbes to intercept sources they cannot intercept under purely attractive chemotaxis

We next considered how a microbe might intercept a moving source more efficiently than is allowed by simply following the gradient of an attractant. In particular, we explored how signals from two chemoeffectors might be combined as there will typically be multiple nutrients being simultaneously released by any real source (Fig 2A). We noted that when two nutrients being released by a moving source have different diffusion coefficients the gradients of their concentrations will point in different directions (Fig. 2B), whereas if the source were stationary the gradients would have different magnitudes but point in the same direction. Because the gradients point in different directions a weighted sum or difference of the two gradients could be constructed to point in any desired direction within the relevant 2D cross-section (Fig. 3C). While both nutrients are primarily concentrated in a plume behind the source, the slow-diffusing nutrient trails behind the source more than the fast-diffusing nutrient (Fig. 2B). Responding to the fast-diffusing nutrient as an attractant and the slow-diffusing nutrient as a repellent could therefore be a potential means of successfully aiming in from of the moving source.

**Fig. 2.**
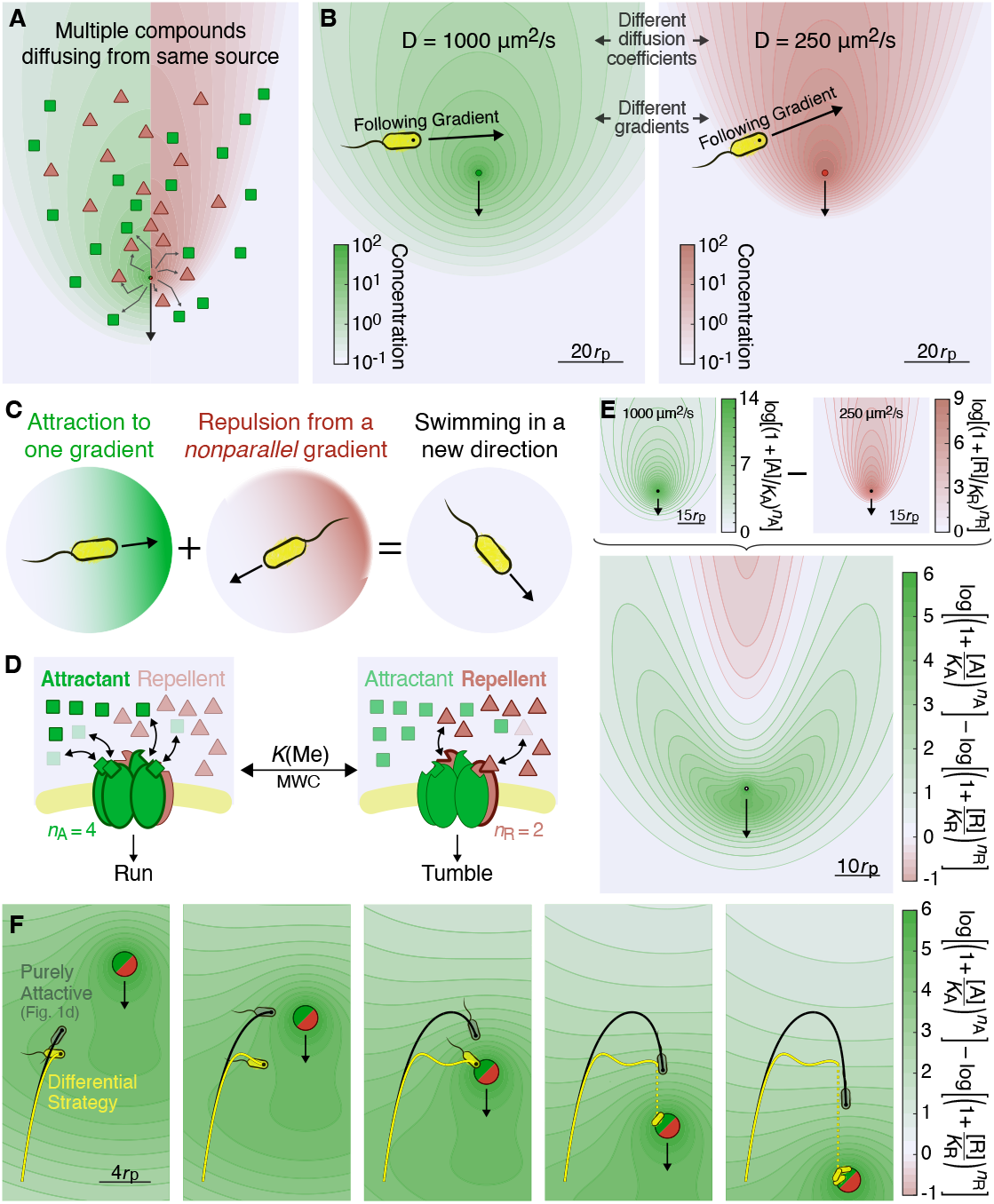
Potential for the differential use of two concentration gradients to improve chemotaxis towards a moving source. **A** Cartoon illustrating a moving source of multiple nutrients. **B** Comparison of the nutrient plumes created by two nutrients with different diffusion coefficients being released at 1 fmol/s from the same moving source. Source is a particle of radius 10 μm moving at 9 μm/s. Concentration is normalized to the microbe’s sensitivity to each nutrient (100 nM). **C** Visual representation of the core concept behind differential chemotaxis: when two gradients are nonparallel, following a sum, difference, or other function of the two can allow a microbe to swim in any direction in the relevant plane. **D** Simplified representation of a heterogenous Monod-Wyman-Changeux model for *E. coli*’s chemotactic response. A cluster of *n*_*A*_ = 4 attractant receptors and *n*_R_ = 2 repellent receptors are either all in an inactive state that promotes continued running or all in an active that promotes tumbling. Attractants and repellents can only bind the inactive and active states, respectively. Adaption based on receptor methylation ensures the response depends on gradients rather than absolute concentrations (Methods). **E** Illustration of how the attractive (top left) and repulsive (top right) components of the differential strategy combine to create a differential response function (bottom) whose gradient the microbe can follow to correctly aim in front of the source. The attractant and repellent are each being released at 1 fmol/s and the microbe’s receptor binding affinities (*K*_*A*_ and *K*_R_) are both 100 nM. **F** Trajectory for a microbe starting from the same location as in Fig. 1D but now following the differential strategy (yellow line and microbe) compared to the trajectory from Fig. 1D (black line and gray-green microbe). Frames show evenly spaced timepoints from 25.7 to 44.3 seconds. The microbe following the differential strategy successfully intercepts the particle while the microbe following the purely attractive strategy does not. In our simulations, the particle is intercepted when the point the microbe is located at (black dot) reaches the surface of the particle (without consideration of the size of the microbe, which is not drawn to scale in this nor any other figures).

**Fig. 3.**
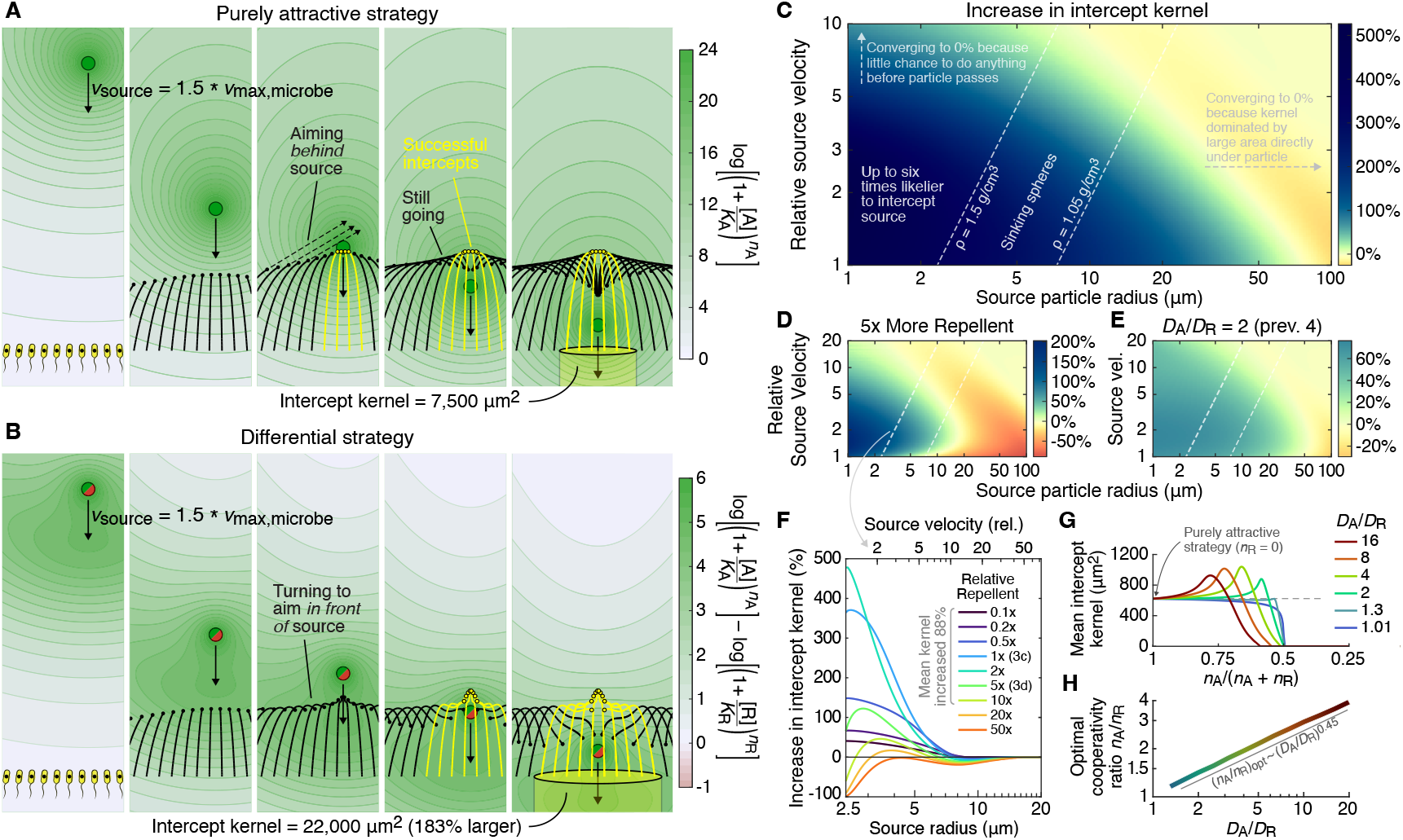
The differential strategy significantly increases the likelihood of intercepting a source moving faster than the microbe can swim and does so over a wide range of conditions. **A** Set of trajectories illustrating the “intercept kernel” for a microbe following a purely attractive chemotactic strategy. Microbes’ current locations are represented by black dots and their trajectories by black lines. After microbes intercept the source particle their trajectories are colored yellow. Microbes starting up to 49 μm away from the axis the source is traveling along successfully intercept the source particle, which corresponds to a 7,500 μm^2^ disk of initial positions from which microbes will successfully intercept the moving source. This disk is the intercept kernel and is proportional to the probability of a microbe with a random initial position successfully intercepting the moving source. Same parameters as in Fig. 2E-F. Second through fifth frames show timepoints evenly spaced from 22.5 to 40.5 seconds after the first frame. **B** Same as **A** but now showing trajectories for microbes following the differential strategy. Microbes following the differential strategy are able to intercept the particle from initial positions 68% further from the source’s path than microbes following a purely attractive strategy, resulting in a 183% larger intercept kernel (meaning a microbe with a random initial position would be 2.83 times more likely to intercept the source). By the third frame microbes following the differential strategy are consistently aiming in front of the source, which is the key to the increase in intercept kernel. **C** Increase in intercept kernel for the differential strategy compared to the purely attractive strategy for a range of source particle radii and velocities (normalized to the microbe’s maximum velocity). Diagonal white dashed lines show the approximate velocities at which a spherical particles with densities of 1.5 g/cm^3^ and 1.05 g/cm^3^ would sink (Methods). **D** Same as **C** but now with the repellent being released from the source at five times the rate as the attractant (5 fmol/s, instead of 1 fmol/s). For some source particle characteristics the differential strategy decreases the intercept kernel, but for particles sinking at realistic speeds given their radii (same white dashed lines as in **C**) only a small decrease is observed (as is further explored in **F**). **E** Same as **C** but now with less difference between the diffusion coefficients of the attractant and repellent (*D*_*A*_/*D*_R_ = 2, whereas 4 was used in previous figures and calculations). The average numbers of attractant and repellent receptors per cluster have been adjusted to *n*_*A*_ = 3.6 and *n*_R_ = 2.4 to account for the dependence of their optimal ratio on the diffusion coefficient ratio (as explored in **G** and **H**). **F** Increase in intercept kernel for the differential strategy as compared to the purely attractive strategy as a function of source particle radius (x-axis) for different rates of repellent release (colormap, defined relative to the attractant release rate, which is held constant at 1 fmol/s). Source velocity is calculated based on source particle radius and assuming spherical particles of density 1.5 g/cm^3^ sinking through water (leftmost of the white dashed lines in **C-E**). For source radii log-uniformly distributed from 2.35 to 20 μm and relative repellent release rates log-uniformly distributed from 0.1 to 10, the mean intercept kernel for the differential strategy is 88% larger than the mean kernel for the purely attractive strategy – which means a microbe following the differential strategy is nearly twice as likely to intercept a moving source whose characteristics are sampled from these distributions than a microbe following the purely attractive strategy. **G** Geometric mean intercept kernel as a function of the fraction of attractant receptors in each cluster for different diffusion coefficient ratios taken across source radii log-normally distributed from 2.5 to 20 μm with source velocities calculated for sinking spheres of density 1.5 g/cm3. **H** Cooperativity ratio *n*_*A*_/*n*_R_ that corresponds to the largest intercept kernel (calculated as in **G**) as a function of the diffusion coefficient ratio *D*_*A*_/*D*_R_.

To investigate how specifically a microbe should respond to the combination of a fast-diffusing attractant and slow-diffusing repellent, we expanded our model of chemotactic receptor binding into a heterogenous Monod-Wyman-Changeux model^53^ based on detailed modeling of *E. coli* chemotaxis by Mello and Tu 2005^55^, Park and Aminzare 2021^26^, and others^10,31,46,53,54^ with some additional simplifications (Methods). In our model, a mix of attractant and repellent receptors form a cluster that is either entirely in a run-promoting inactive state or entirely in a tumble-promoting active state (Fig. 3D, Methods). The attractant only binds its receptor in the inactive state and the repellent only binds its receptor in the active state (Fig. 3D). With receptor methylation acting to maintain an equal ratio of active to inactive clusters (Methods, Supplemental Information), this model predicts the microbe should follow the gradient of the “differential response function”

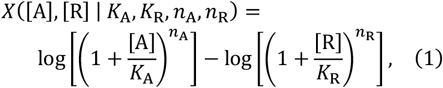

where [A] and [R] are the attractant and repellent concentrations, *K*_*A*_ = *K*_R_ = 100 nM are the attractant and repellent binding affinities, and *n*_*A*_ = 4 and *n*_R_ = 2 are the average number of attractant and repellent receptors in each cluster (with *n*_*A*_ + *n*_R_ = 6 chosen based on *E. coli*’s canonical trimer-of-dimers arrangement and *n*_*A*_/*n*_R_ = 2 chosen for being the roughly optimal ratio based on results presented later in this paper). Unlike purely attractive chemotaxis (e.g. the *n*_R_ = 0 case of Eqn. 1), the differential response function has larger values just in front of the particle than behind it (Fig. 2E). A microbe with receptors for a fast-diffusing attractant and a slow-diffusing repellent in heterogeneous clusters would therefore aim in front of (rather than behind) a moving source and may be able to intercept sources a microbe with only attractant receptors could not.

To test whether this distinct change in trajectory would indeed translate into successful intercepts, we repeated the previous simulations (Fig. 1D, in which the microbe had followed a “purely attractive” strategy defined by the *n*_*A*_ = 6 and *n*_R_ = 0 case of Eqn. 1) but now for a microbe following a “differential” strategy (with *n*_*A*_ = 4 and *n*_R_ = 2 as described above) and compared the two trajectories (Fig. 2F). Initially, both microbes swam upwards as only the faster-diffusing attractant had a significant concentration while the source was still relatively far away. But, as the source got closer, the microbe following the differential strategy had a sudden downwards turn while the purely attractive microbe continued to swim upwards (Fig. 2F). This downwards turn put the differential microbe on a nearly optimal intercept trajectory and allowed it to successfully intercept the moving source while the purely attractive microbe failed to ever intercept it (Fig. 2F). Thus, a differential chemotactic strategy can allow a microbe to break out of the inherent limitations of following concentration gradients to reach a moving source and successfully intercept moving sources that microbes following purely attractive chemotactic strategies cannot.

### A differential strategy robustly increases the likelihood of intercepting a moving source as compared to a purely attractive strategy

Having demonstrated that combining attraction to a fast-diffusing nutrient and repulsion from a slow diffusing nutrient can allow a microbe to successfully intercept a moving source that it could not when only responding to the attractant, we proceeded to test whether this differential strategy would be robustly beneficial over a range of conditions. To quantify performance, we defined the “intercept kernel” as the cross-sectional area of the range of initial positions from which a microbe could successfully intercept the source. Specifically, we found the maximum radial distance from the axis the source is traveling along that a microbe could start from and still intercept the particle (Methods) and calculated the intercept kernel as the area of the corresponding disk of initial positions (Fig. 3A). This intercept kernel is directly proportional to the probability of a microbe with a random initial position intercepting the source. We therefore compared the intercept kernels under the differential and purely attractive strategies to quantify how much more likely a microbe following the differential strategy would be to successfully intercept the source particle. For example, for the particle radius and velocity used previously (10 μm and 9 μm/s), the intercept kernel is 183% larger under the differential strategy (Fig. 3B vs Fig. 3A), which means microbes would be 2.83 times likelier to successfully intercept the source if following the differential strategy. Across source radii from 1 to 100 μm and velocities from 1 to 10 times the microbe velocity, the differential strategy increases the intercept kernel by 117% on average (meaning microbes would be more than twice as likely to intercept the source) with a more than 6-fold increase for small sources moving about twice as fast as the microbe (a parameter regime that could be relevant to predators intercepting their prey) (Fig. 3C). Thus, employing a differential strategy greatly increases the likelihood of a microbe successfully intercepting a moving source.

We next tested the differential strategy against what should be its biggest vulnerability: an increase in the amount of repellent released by the moving source. Intuitively, being repelled by anything released from the source carries the risk of that repulsion outweighing any attraction and the microbe actively swimming away. There will inevitably be some but also some release rate that is too small to significantly impact the trajectories. A microbe could in principle evolve appropriate receptor binding affinities to each nutrient to keep the expected release rates (as normalized to its binding affinities) in the range in which a differential strategy would be beneficial. But, if the variation in the nutrients’ release rates between source particles were larger than the differential strategy is robust to then the differential strategy could cease to be beneficial on average regardless of receptor binding affinities. To explore how much variation the differential strategy can withstand, we recalculated the increase in intercept kernel over the previous grid of particle radii and velocities but now with five times more repellent than attractant being released by the source (Fig. 3D). Depending on particle characteristics, the differential strategy could now be either very beneficial (increasing the intercept kernel more than 3-fold) and very detrimental (decreasing the kernel nearly 10-fold) (Fig. 3D). We noted, however, that the largest decreases in intercept kernel occurred for particles sinking abnormally slowly for their size, which would minimize the consequence of this region in realistic scenarios. We therefore narrowed our focus to particles with radii between 2.5 and 20 μm and velocities calculated for sinking spheres of density 1.5 g/cm^3^ (Methods) before exploring a wider range of relative repellent release rates (Fig. 2F). As expected, at the highest repellent release rates the differential strategy never outperformed the purely attractive strategy. However, over a broad range of repellent release rates from 0.1 to 10 times the attractant release rate and across the array of source particle radii, the differential strategy generally outperformed the purely attractive strategy (Fig. 2F) with the average intercept kernel under the differential strategy being 88% larger than under the purely attractive strategy. Additionally averaging over densities between 1.05 and 1.5 g/cm^3^ also continued to yield an improvement in the intercept kernel (78%, Appendix Fig. S4, Methods). Thus, even if the relative release rates of the attractant and repellent vary across two orders of magnitude, a microbe could receive a sizeable average benefit from employing a differential strategy – illustrating a surprising robustness against what could have been a critical vulnerability.

The final source of natural variation that we considered was the microbe’s own relative expression of attractant and repellent receptors. As we performed this analysis we also explored the effectiveness of the differential strategy under different ratios of attractant and repellent diffusion coefficients. We began by calculating the geometric mean intercept kernel as a function of the average ratio of attractant to repellent receptors in each cluster (Methods) for different diffusion coefficients (Fig. 3G). We observed a differential strategy to be beneficial for any ratio of attractant to repellent diffusion coefficients greater than *D*_*A*_/*D*_R_ ≈ 1.3 provided the average number of attractant receptors per cluster was sufficiently large (Fig. 3G). It was typically possible to decrease the fraction of attractant receptors in each cluster by about 9% from the optimum composition or increase it by any amount with the differential strategy still being beneficial (Fig. 3G). We also observed the optimal ratio of attractant to repellent receptors to scale as approximately the square root of the ratio of the diffusion coefficients (specifically as *n*_*A*_/*n*_R_ = (*D*_*A*_/*D*_R_)^0.45^, Fig. 3H). Adjusting the average cluster composition to *n*_*A*_ = 3.6 and *n*_R_ = 2.4 based on these results, we calculated the increase in the intercept kernel across the grid of source particle radii and relative velocities with the diffusion coefficient of the repellent now set to 500 μm^2^/s. Across the vast majority of this parameter range (approximately three quarters) the differential strategy was indeed beneficial and increased to intercept kernel by up to 75% (Fig. 3E). Thus, the differential strategy is robust to variations in the microbe’s relative expression of attractant and repellent receptors and can be employed for combinations of attractants and repellents with a wide range of diffusion coefficients.

### A differential strategy continues to increase the likelihood of intercepting a moving source when explicitly modeling stochastic run-and-tumble motion

In all results presented thus far, we have worked with a deterministic approximation in which microbes swim exactly in the direction of the nutrient concentration or differential response function gradient with a velocity based on averaging over a large series of runs and tumbles (Methods). Bacterial chemotaxis is, however, a sufficiently stochastic process that the uncertainty in the microbe’s position is not necessary negligible compared to its expected motion. For example, if a microbe like *E. coli* swims at 30 μm/s with an average run length of 1 s and no correlations between run direction, its diffusion coefficient would be approximately 900 μm^2^/s and the typical uncertainty in its position by the time it intercepts the particle in our simulations would be more than 100 μm, which is considerable larger than most of the particle radii we explored. Therefore, to confirm that differential chemotaxis is still a beneficial strategy even with the full stochasticity of bacteria chemotaxis considered, we ran explicit run-and-tumble simulations in which the microbe “ran” in a straight line at constant velocity and then “tumbled” to a new random orientation at a frequency that depended on the rate at which it observed the response function to change (Methods). Parameters were guided by experimental observations^11,45,46^ and chosen to match the expected net velocity (from averaging over all possible runs) to what had been used in the deterministic approximation (Methods). Three sets of parameters provided three levels of stochasticity (Methods). Depending on the level of stochasticity, the differential strategy increased the intercept kernel between 5 and 16% (Supplementary Fig. S2). This improvement is notably smaller than the improvement obtained under the deterministic approximation (a 98% increase, or nearly double, for the same source particle characteristics) and is brought down by microbes now being able to “get lucky” regardless of strategy (with even randomly motile, non-chemotactic bacteria now having a small rate of intercepting the particle) or “get unlucky” and never come out of a tumble facing in the correct direction for the final intercept (a distinct possibility when only ∼10% of possible orientations yield a successful intercept when the microbe is 1 run length away). Nevertheless, when successfully intercepting a food particle can make the difference between proliferation and starvation, even small increases in the intercept kernel can represent consequential fitness differences. Thus, differential chemotaxis remains a potentially highly beneficial strategy even when incorporating the inherent stochasticity of bacterial chemotaxis.

### A differential strategy is also beneficial to faster-swimming run-and-reverse marine microbes

As a final test of the differential chemotaxis strategy, we adapted our modeling to now reflect a marine microbe attempting to intercept small sinking particulate organic matter (whereas *E. coli* had guided choices thus far). Of great relevance to adapting our simulations is that marine microbes generally swim much faster than *E. coli*^36,47,57–60^, which also increases the relevant range of source velocities to use. Faster source particles raised the Péclet numbers enough to motivate numerically calculating concentration plumes using finite element analysis (Methods). Recent work by Słomka et al. 2020^39^ has also shown that fluid flows around sinking marine particles significantly impact the likelihood of marine microbes intercepting these particles. For our marine microbe simulations we incorporated both the translational and rotational effects of fluid flows on the microbe trajectories following work by Słomka et al. 2020^39^ (Methods). As marine microbes frequently employ run-and-reverse or run-and-flick dynamics instead the run-and-tumble of *E. coli*^47,61,62^, we implemented stochastic run-and-reverse dynamics in which a microbe swam in a straight line until reversing its direction with a probability that depended on local nutrient concentrations while keeping the same differential response function for calculating the frequency of the microbe reversing (Methods). Because the velocities of both the microbe and source particle were increased, the relevant timescales shrunk sufficiently that microbes were likely to only perform a couple reverses before the source had passed, making a deterministic approximation based on averaging over many runs inappropriate and leading us to only perform a stochastic simulation (Methods). Using these modeling modifications and with microbe and source velocities of 150 μm/s and 200 μm/s (much faster than in previous simulations), we observed the differential strategy to increase the probabilistic intercept kernel by approximately 6% (Supplementary Fig. S3). As these marine-microbe simulations and the previous simulations guided by *E. coli* chemotaxis represent two very different scenarios, these results suggest differential chemotaxis should be beneficial to a diverse range of microbes in widely varying conditions.

### E. coli is attractant to fast-diffusing amino acids and repelled by slow-diffusing amino acids, as is necessary to implement the differential strategy

Having shown the theoretical potential of a differential strategy, we proceed to establish a connection to existing experimental data. A thorough characterization of which amino acids are attractants and repellants for *E. coli* was performed by Yang et al. 2015^18^. We used characterizations of MG1655 *E. coli* from Yang et al. 2015^18^ and compared to diffusion coefficients for each amino acid (Fig. 4). As reported diffusion coefficients vary between sources, we assembled diffusion coefficients from several sources^63–68^ and kept the reported values separated by source with no adjustments to the values to correct for measurement differences. A striking trend emerged in which all 8 attractants had diffusion coefficients greater than 790 μm^2^/s in all reported measurements and all 7 repellants had diffusion coefficients less than this cutoff in all reported measurements except for two outliers (Fig. 4). This attraction to fast-diffusing amino acids and repulsion from slow-diffusing amino acids is exactly what would be needed to implement the differential strategy. Thus, in addition to the theoretical benefits of differential chemotaxis, the strong correlation between the amino acid diffusion coefficients and *E. coli*’s chemotactic response to them provides evidence that microbes may actually implement this strategy.

**Fig. 4.**
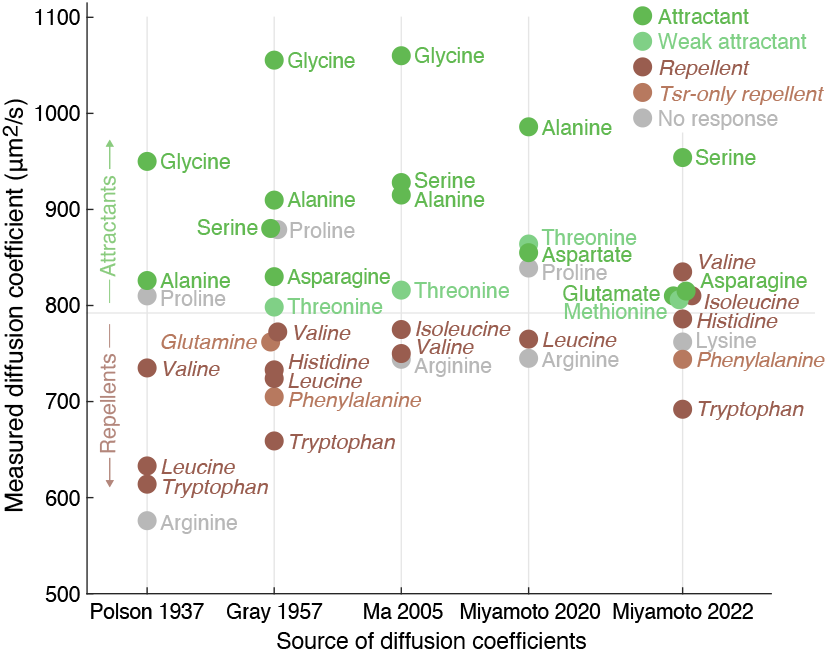
E. coli is attracted to fast-diffusing amino acids and repelled by slow-diffusing ones. Diffusion coefficients of the amino acids (vertical axis) along with *E. coli*’s chemotactic response as reported by Yang et al. 2015^18^ (colors) separated by source^63–68^ (horizontal axis). Amino acids that were described by Yang et al. as weak attractants are colored separately as are amino acids that were only repellents to the Tsr-only *E. coli* strain. Diffusion coefficients for glycine, asparagine, glutamine, proline, histidine, phenylalanine, and tryptophan in the “Gray 1957” column were originally measured by Longsworth 1953^68^ before being included in Gray’s table. Horizontal line separating attractants from repellents is an empirical observation based on data.

## Discussion

In this paper we demonstrated that a differential strategy combining attraction to a fast-diffusing nutrient with repulsion from a slow-diffusing nutrient can significantly improve chemotaxis towards a moving source. Employing this strategy can increase the likelihood of intercepting a fast-moving source more than six-fold and is robustly beneficial across a range of initial conditions and microbe and source particle characteristics (Fig. 3). We also observed that *E. coli* is attracted to the fast-diffusing amino acids but repulsed by the slow diffusing ones (Fig. 4). There are other convincing explanations for why *E. coli* would be repulsed by certain amino acids, including the correlations to utility as a nutrient and growth inhibition explored in Yang et al. 2015^18^, but ultimately multiple explanations for this behavior can be true as traits can be beneficial for multiple reasons. Additionally, the specific example of repulsion from tryptophan is intriguing as it does not inhibit growth even at high concentrations^18^ and can be utilized as *E. coli’s* sole carbon and nitrogen source^18^. However, tryptophan is the slowest diffusing amino acid, which makes repulsion from it especially beneficial under the differential chemotaxis explanation. The benefits of differential chemotaxis may therefore help explain some of the mystery behind *E. coli’s* repulsion from certain amino acids.

We considered deterministic trajectories throughout most of the Main Text, but microbial motion is stochastic, so we extended our results with stochastic modeling in the Supplementary Information. In particular, microbes do not perfectly follow gradients but instead display run-and-tumble motion in which they “run” in a roughly straight line and then randomize their orientation by tumbling in response to worsening conditions before starting another run^11^. Stochastic run-and-tumble simulations with parameters reflecting *E. coli* chemotaxis produced increases to the probabilistic intercept kernel from employing the differential strategy between 5 and 16% (Methods, Supplementary Fig. S2), while stochastic run-and-reverse simulations of marine microbes also showed an approximately 6% increase (Methods, Supplementary Fig. S3). These increases are smaller than what was observed for the deterministic simulations due to microbes being able to get “lucky” or “unlucky” regardless of strategy. Success or failure to intercept a food particle does, however, have huge fitness consequences (potentially the difference between proliferation and starvation), so a 5 to 16% increase in the likelihood of doing so is significant. Thus, while additional work can confirm the breadth of applicability under variable conditions and assumptions, the benefits of differential chemotaxis appear robust to the inherent stochasticity of microbial chemotaxis.

The simulations presented in the Main Text used a point-source calculation of concentration plumes and ignored the effects of fluid flows around the source particle, so it is important to determine in which regimes these calculations are valid. Kiørboe et al. 2001^69^ showed that if the particle is large and fast-moving (i.e. has a large Péclet number) then the nutrient concentration plume becomes long and narrow and no longer resembles the plumes we calculated. Słomka 2020^39^ meanwhile provides a highly detailed account of how fluid flows around a particle impact microbe trajectories through both translational and rotational effects. To confirm that neither the point-source approximation for calculating concentration plumes nor the neglecting of fluid flows around the particle significantly impacted our results, we repeated our simulations with these effects now accounted for (Supplementary Fig. S1). When using the numerically calculated concentration plumes intercept kernels changed by just 15% on average (mean of the absolute value of the change from kernels calculated using the point source approximation) and the average increase in the intercept kernel remained at 95% (Supplementary Fig. S1). Accounting for the advective effects of fluid flows on microbe trajectories had a larger impact on our results and for large particles worsened the performance of the differential strategy (Supplementary Fig. S1), but for small particles with both sets of effects accounted for increases in the intercept kernel of up to 254% (meaning microbes would be 3.5 times likely to successfully intercept the source particle) were still observed (Supplementary Fig. S1). The simulations of marine microbes described in the Results and Supplementary Fig. S3 also used numerically calculated concentration plumes, accounted for both the translational and rotational effects of fluid flows on microbe trajectories, and observed the differential strategy to still increase the probabilistic intercept kernel (Methods, Supplementary Fig. S3). Thus, our results do not appear to be critically dependent on the simplifications we made, and differential chemotaxis can be a beneficial strategy even when considering the complicated effects of fluid flows around the source.

An interesting property of the differential strategy is that, although it increases the likelihood of a microbe successfully intercepting a moving source, when a microbe can intercept the source under either a differential or purely attractive strategy the microbe usually takes longer to reach the source under the differential strategy (e.g. Fig. 3A-B). This increase in the time to reach the source occurs because the microbe’s decreased investment in attractant receptors and the attractive and repulsive components of the differential response function partially canceling each other mean the microbe is following a less steep gradient and therefore has a slower net velocity. If, however, the microbe’s velocity is constant regardless of the magnitude of the response function gradient, then the differential strategy will actually consistently decrease rather increase the time to reach the source, as we previously explored^70^. Additionally, the increase in time to the reach the source is limited to only a couple seconds due to the short time scales over which particle intercepts happen. Thus, compared to fitness benefit of either successfully intercepting a particle or failing to do so, the impact of strategy choice on the time it takes a microbe to reach the source is largely inconsequential.

While we considered a sinking particle throughout this paper, a differential strategy may be the most relevant to microbes attempting to intercept or follow other microbes for the purpose of predation or crossfeeding. This relevance is due to the differential strategy being most beneficial when the source is small (e.g. around the size of a cell) and moving about twice as fast as the chemotactic microbe’s net velocity. The latter condition is satisfied if a predator microbe’s straight-line swim speed matches that of its prey and it navigates with a chemotactic efficiency (i.e. the ratio of expected net velocity vs straight-line speed) of 50%^47^. In the context of crossfeeding, marine bacteria have been observed to successfully track photosynthetic algae, presumably for the purposes of feeding off leaked and exuded nutrients^42^, despite some modeling suggesting the information in these nutrient plumes should not be sufficient to allow successful tracking^47,71–73^. In part because the relevant success metrics (either integrated exposure to the nutrient plume or the expected time the algae can be tracked before it is lost) are so different than simply intercepting a source particle, exploring the benefits of differential chemotaxis to microbe attempting to track or follow other microbe was beyond the scope of this paper. Future theoretical and experimental research should therefore explore how the marine microbes that associate with algae respond to the different components of the nutrient plume and whether a differential strategy contributes to their ability to successfully track their targets.

Another important direction for future work will be to experimentally characterize trajectories. A variation on the microfluidic chip used in Stocker et al. 2008^36^ (in which an attractant was injected upstream of a circular pillar surrounded by a fluid flow to simulate release from a sinking particle) might provide a convincing experimental demonstration. The first step towards an experimental demonstration may, however, be to simply place bacteria in orthogonal attractant and repellent gradients and see if they actually do swim on “diagonal” trajectories. Future experimental and theoretical work can clarify how microbes actually combine the signals of attractants and repellants to optimize chemotaxis in natural, dynamic environments, as it appears from our work that attraction and repulsion have the potential to combine in surprising and remarkably beneficial ways. Indeed, while we presented the benefits of a differential strategy specifically to intercepting a moving, we hope the more foundational observation that integrating information from multiple concentration gradients can produce chemotaxis in otherwise inaccessible directions inspires a variety of future research directions and unlocks insights into how microbes navigate the various spatially complex environments they inhabit.

## Methods

### Nutrient concentrations from point source approximation

Most simulations used a point-source approximation for the nutrient concentrations, in which concentrations were calculated by integrating over the concentration plumes for the instantaneous nutrient release time *t* ago according to

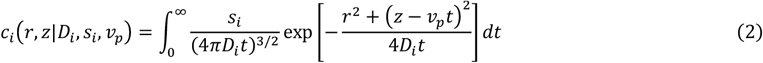

where *r* and *z* are cylindrical coordinates for a system centered on the source’s current location, *D*_*i*_ is the nutrient’s diffusion coefficient, *s*_*i*_ is the rate at which it is being released from the source, and *v*_*p*_ is the source particle velocity (downwards, in the 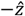 direction). The nutrient plume is calculated for diffusion in 3D space, but is azimuthally symmetric under rotations around the axis the source is traveling along, so only depends on *r* and *z* but not *ϕ*. Performing this integration yields

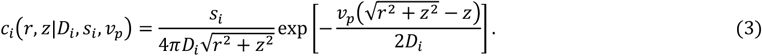

In addition to treating all nutrient as being released from the center of the source particle, the point source approximation also neglects any fluid flows around the particle and instead assumes uniform, symmetric diffusion.

In most simulations (except Fig. 3E, G, and H, in which different diffusion coefficients are being actively discussed) the attractant diffusion coefficient is *D*_*A*_ = 1000 μm^2^/s and the repellent diffusion coefficient is *D*_R_ = 250 μm^2^/s. The attractant and repellent release rates are *s*_*A*_ = *s*_R_ = 1 fmol/s (except in Fig. 3D and F, in which the repellent release rate is varied).

### Nutrient concentration form finite element analysis

The point source approximation is valid when the Péclet number (defined as 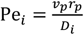 where *r*_*p*_ is the source particle radius and representing the ratio of the importance of advection and diffusion) is less than ∼1, as is the case in the majority of simulations performed in this paper. (Specifically, at small Péclet numbers, the concentration plume calculated with the point-source approximation roughly matches the numerical calculation of the concentration plume accounting for finite-size of the source; Supplementary Fig. S1.) The phase spaces presented in Fig. 3C-E do, however, include sufficiently large source particle radii and velocities in their top-right corners to raise the Péclet numbers above 1. For this reason, the phase space in Fig. 3C is reproduced in the Supplementary Fig. S1 using numerically calculated concentration plumes. The simulations of marine microbes also used numerically calculated concentration plumes.

The numerically calculated concentration plumes where calculated using finite element analysis in Mathematica (using the `NDSolve` function with the `Method -> {“PDEDiscretization” -> “FiniteElement”}` option) to solve the steady-state advection-diffusion equation

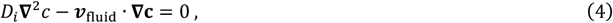

or in cylindrical coordinates with no azimuthal dependence

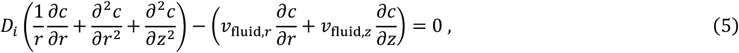

with a Dirichlet boundary condition (specifically a constant concentration across the source particle surface) and Stokes flow around a spherical particle given by

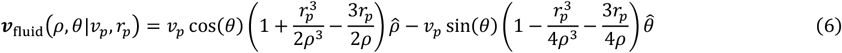

where ***v***_fluid_ is the fluid flow in the moving frame of the source particle and *ρ* and *θ* are spherical coordinates centered on the source particle with *θ* = 0 corresponding to up (i.e. 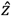 in the cylindrical coordinate system). After performing the finite element analysis the nutrient flux *J* through the surface of the spherical source particle was calculated from the derivative of the concentration according to

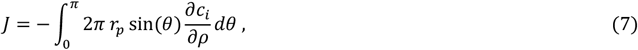

and the concentration plume was scaled by a constant factor of *s*_*i*_/*J* to obtain the desired nutrient release rate.

### Optimal trajectories as presented in Fig. 1B and 1C

The optimal trajectory is the trajectory which, for a fixed microbe swim speed (6 μm/s in Fig. 1B and 1C), would result in the microbe intercepting the particle at the earliest possible time. It is a straight line trajectory and we define it by the angle *θ*_opt_ formed between the trajectory and the vector pointing directly from the microbe to the particle’s current location. Intuitively, it is how far in front of the particle the microbe should ideally aim.

The optimal trajectory can be determined by the law of sines. A triangle is formed by the point **P** where the source particle starts, the point **M** where the microbe starts and the point **I** where the microbe intercepts the particle. The microbe’s initial position relative to the particle defines the angle *θ*_0_ at vertex **P** and we define the optimal trajectory by the angle *θ*_opt_ at vertex **M**. The law of sines states that

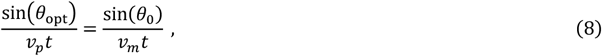

where *v*_*p*_ and *v*_*m*_ are the particle and microbe velocities and *t* is the time until the intercepts the source particle. The integral

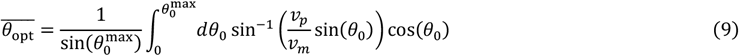

can then be solved analytically to yield the mean optimal trajectory in a spherical conical region of half-width 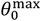, and a similar integral can be solved to yield the standard deviation of the optimal trajectory in that region.

### Chemotactic response and microbe velocity

As noted in the Results, the differential response function is given by

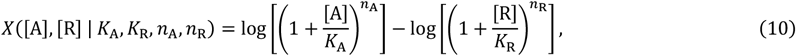

where [*A*] = *c*_*A*_(*r, z*) and [R] = *c*_*R*_(*r, z*) are the local concentrations of the attractant and repellent, *K*_*A*_ and *K*_R_ are the receptor binding affinities to each, and *n*_*A*_ and *n*_R_ are the average number of attractant and repellent receptors in each cooperative cluster. The purely attractive strategy (including as used in Fig. 1D) is the {*n*_*A*_, *n*_R_} = {6,0} case of Eqn. 10. This response function is derived in the Supplementary Information and can also be obtained by simplifying results in Mello and Tu 2005^55^ and Park and Aminzare 2021^26^ under an appropriate assumption about the timescale on which receptor methylation occurs (Supplementary Information).

In the deterministic approximation, the microbe’s velocity at small values of |∇*X*| was calculated from Equation 3.1 of Colin et al. 2014^46^ and determined to be

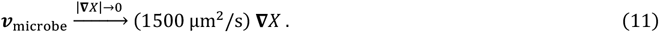

We additionally imposed a maximum velocity of 6 μm/s based on experimental observations^45,46^ to arrive at

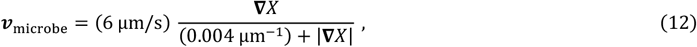

which was used in all deterministic approximations in this paper.

All calculations and simulations assume 3D space, but because the nutrient plume has an azimuthal symmetry about the vertical axis to the source is traveling along all motion occurs in a 2D plane that includes this axis and the microbe’s initial position.

### Calculating intercept kernels

In deterministic simulations, intercept kernels were calculated by finding the highest point on the source particle which absorbed microbes (i.e. satisfying 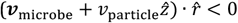) and integrating *v*_microbe_ in reverse until the trajectory was far enough below the particle that any additional radial motion was insignificant. In the Supplementary Fig. S1 simulations in which the fluid flows around the source particle were considered, the topmost point on the particle often absorbed microbes, so when this was the case we initiated the backwards integration from the highest point directly above the particle which had a downwards flux of microbes (in the particle frame). Occasionally, the backwards integration of a trajectory reach a limit cycle or fixed point in the particle frame or intersected the particle at a lower location, so when this was the case we stepped the trajectory termination point down the particle until a feasible trajectory was obtained.

In stochastic simulations, intercept kernels were calculated slightly differently for the *E. coli* run-and-tumble simulations and the marine microbe run-and-reverse simulations. Calculation of those intercept kernels is detailed at the ends of each of the two corresponding sections below.

### *E. coli* run-and-tumble simulations

For the run-and-tumble simulations we used a simplified model, formulated to facilitate matching of the expected drift velocity to the deterministic approximation used in the Main Text while accurately reflecting experimentally established models of *E. coli* chemotaxis. Specifically, in our simulations the microbe “ran” by swimming in a straight line at velocity *v*_microbe_ and “tumbled” at a probability

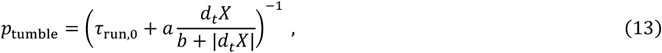

where *d*_*t*_*X* is the temporal derivative of the response function as experienced by the microbe, the mean run length *τ*_run,0_ = 1 s in all simulations, and *a* and *b* were varied as described below. “Tumbles” lasted 150 ms during which the microbe did not swim in any direction and after which it had a new orientation, which was uniform-randomly sampled in 3D space.

Three microbe velocities were used, and for each velocity the parameters *a* and *b* were set such that the microbe’s expected net velocity obtained by averaging over the expected distance traveled on all possible runs divided by the combined run and tumble time according to

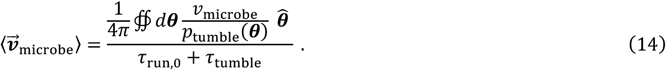

The {*v*_microbe_, *a, b*} combinations used are provided in Table 1. All combinations produce the same expected velocity according to Eqn. 14. However, an approximate diffusivity can also be calculated as 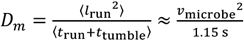, which highlights each parameter set as representing a different level of stochasticity.

**Table 1.** Parameters for run-and-tumble simulations and corresponding shallow-gradient velocities, steep-gradient velocities, and microbe diffusivities obtained from averaging over runs in all position directions.

| $v_{\text{microbe}}$ | $a$ | $b$ | $\left\vert \lim_{\vert d_t X\vert \rightarrow 0} \langle \vec{v}_{\text{microbe}} \rangle \right\vert / \vert \nabla X\vert $ | $\left\vert \lim_{\vert d_t X\vert \rightarrow \infty} \langle \vec{v}_{\text{microbe}} \rangle \right\vert $ | Approximate diffusivity |
| --- | --- | --- | --- | --- | --- |
| 30 $\mu\text{m}^2/\text{s}$ | 0.460 s | 0.080 $\text{s}^{-1}$ | 1500 $\mu\text{m}^2/\text{s}$ | 6 $\mu\text{m}/\text{s}$ | 782 $\mu\text{m}^2/\text{s}$ |
| 24 $\mu\text{m}^2/\text{s}$ | 0.575 s | 0.064 $\text{s}^{-1}$ | 1500 $\mu\text{m}^2/\text{s}$ | 6 $\mu\text{m}/\text{s}$ | 501 $\mu\text{m}^2/\text{s}$ |
| 15 $\mu\text{m}^2/\text{s}$ | 0.920 s | 0.040 $\text{s}^{-1}$ | 1500 $\mu\text{m}^2/\text{s}$ | 6 $\mu\text{m}/\text{s}$ | 196 $\mu\text{m}^2/\text{s}$ |
| Deterministic model | | | 1500 $\mu\text{m}^2/\text{s}$ | 6 $\mu\text{m}/\text{s}$ | 0 |

Microbes in these simulations were initiated 500 μm below a source particle of radius 20 μm sinking at 9 μm/s, between 0 and 500 μm radially away from the axis the source is traveling along, and with random orientations in 3D space. Their trajectories were simulated until they either intercepted the source particle or fell sufficiently far behind it that they were unlikely to ever catch back up. The intercept kernel was calculated by integrating over the probability of intercepting the source as a function of initial radius weighted by 2*πr*.

### Marine microbe run-and-tumble simulations

For the marine microbe simulations, we simulated a particle of radius *r*_p_ = 30 μm sinking at *v*_p_ = 200 μm. In the particle frame the fluid flow around the particle is given by Eqn. 5 above, and the concentrations of the nutrients were calculated according to the finite element analysis described in the latter part of Methods section “Nutrient concentrations” above and using the same attractant and repellent release rates *s*_*A*_ = *s*_R_ = 1 fmol/s^56^ and diffusion coefficients *D*_*A*_ = 1000 μm^2^/s and *D*_R_ = 250 μm^2^/s.

The microbe has orientation 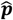, and its position 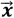, again in the particle frame, evolves according to

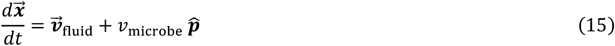

where *v*_microbe_ = 150 μm/s is its swim speed.

The microbe’s orientation 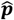 evolves according to Equation 12 in the highly detailed Słomka et al. 2020^39^ under the assumption of a highly elongated microbe (*γ* = 1 in Słomka’s notation). Specifically, 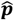 evolves according to

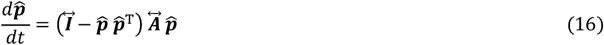

where 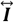 is the identity matrix and 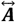 is the velocity gradient defined as *A*_*ij*_ = ∂_*j*_*v*_fluid,*i*_.

Microbes were initiated at 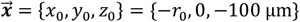 for *r*_0_ from 0 μm to 400 μm and with orientations sampled from a Lebedev Sphere. Microbes swam without reversing for the first 500 ms of the simulations (to prevent “blurring out” of interesting effects related to initial position and orientation) after which they reversed their direction (no more than once due to the short timescale of the simulations) at rate

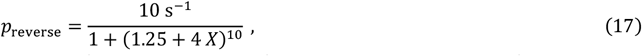

which is tenth-order Hill Function that varies from 10 s^−1^ at *X* → −∞ to 0.1 s^−1^ at *X* → ∞ with a value of 1 s^−1^ at *X* = 0, with *X* being the response function used previously (Eqn. 10 with *n*_*A*_ = 6 and *n*_R_ = 0 for the purely attractive strategy, *n*_*A*_ = 4 and *n*_R_ = 2 for the differential strategy, and *K*_*A*_ = *K*_*R*_ = 100 nM for both strategies). Note that due to the short timescale of these simulations we are neglecting methylation-based adaption such that reversing depending directly on the differential response function *X* rather than on a derivative of it.

Intercept kernels were calculated by running simulations to determine if microbes successfully intercepted the particle and then summing those probabilities over initial positions and orientations weighted by the product of the Lebedev sphere^74^ weights (corresponding to how much area each point is representing, divided by 4π for normalization), the initial radial position (specifically 2*πr*_0_*δr*_0_ to produce an integration over a disk of initial positions), and the normalized upward flux of the microbes in the particle frame (equal to 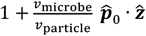 where 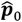 is the microbe’s initial orientation).

In the colormapped plots of intercept probability presented in Supplementary Fig. S3C, the initial orientations used are plotted at evenly spaced orientations on the vertical axis. However, as they come from a 3074-point Lebedev sphere^74^ (which optimizes for even spacing in 3D, and not directly for even spacing in any 2D slice), they are really at positions {0.0°, 3.2°, 8.1°, 13.7°, 19.6°, 25.8°, 32.2°, 38.6°, 45°, 51.4°, 57.8°, 64.2°, 70.4°, 76.3°, 81.9°, 86.8°} + 90° *n* for *n* = {0,1,2,3}. These points differ from even spacing by 2.0 ± 1.1° (mean and standard deviation of absolute value). These plots are however, only for visualization with all reported statistics coming from averaging over all points in the 3D Lebedev sphere with the appropriate weightings.

### Relation between source particle radius and velocity

We used an approximate density and viscosity of water of 1 g/cm^3^ and 10^−2^ cm^2^/s to obtain

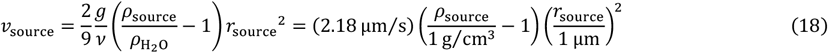

based on Stokes flow around a sphere.

## Supplementary Information *for*

### Derivation of the differential response function from receptor kinetics

This section provides a derivation of the differential response function used throughout the paper from a heterogeneous Monod-Wyman-Changeux (MWC) model of receptor kinetics. The MWC model has a long history of being applied to bacterial chemotactic pathways, and many authors have derived similar results as we present here. (We, in particular, were guided by Mello and Tu 2005^55^ and Park and Aminzare 2021^26^ in developing our model.) Nevertheless, we present a derivation here for convenient reference and to provide clarity on our set of simplifications.

The receptor cluster has *n*_*A*_ attractant receptors and *n*_R_ repellent receptors. The cluster is either entirely in an inactive state that binds the attractant or an active state that binds the repellent. Binding an attractant to its receptor is associated with energy −*k*_B_*T* log ([*A*]/*K*_*A*_), and binding a repellent is associated with energy −*k*_B_*T* log ([R]/*K*_R_). The receptor cluster is methylated by a dynamic adaptation process that creates an energy difference −*k*_B_*T* log(*K*_*M*_) between the active and inactive states. Plugging into a Boltzmann distribution and simplifying yields the probability of the cluster being in the inactive state

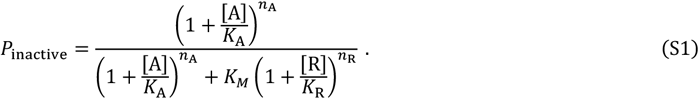

Note that Equation S1 is arrived at because the term 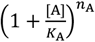 generates 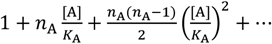 expanded and thus contains the single state of an inactive cluster with no attractant bound, the *n*_*A*_ states with one attractant molecule bound (one state per receptor), and so forth and likewise for the repellent term.)

The temporal derivative of Equation S1 is

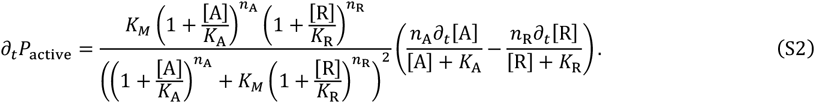

Meanwhile, methylation acting to maintain a 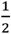 probability of the cluster being active sets the energy difference

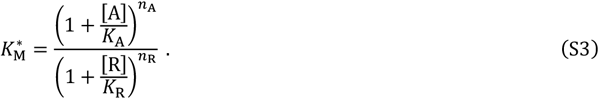

We assume a balance between the rate of change of the chemoeffector concentrations and the rate of methylation such that 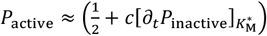.Plugging in Equations S2 and S3 yields

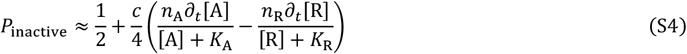

Integrating with respect to time then returns that (up to some constant factors) moving in the direction of the greatest likelihood of the cluster being in the inactive state is the same as following the gradient of

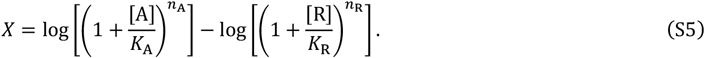

Because Equation S5 is linear in *n*_*A*_ and *n*_R_ (due to the ability to rearrange the exponents out of the logarithms), when the cumulative behavior of multiple clusters of differing receptor composition is considered, *n*_*A*_ and *n*_R_ can simply come to represent the mean number of receptors per cluster instead of the composition of a specific cluster without any change to Equation S5.

We also note that a heterogenous MWC model in which the attractant can bind either of two receptor types (as many amino acids can to both *E. coli* Tsr and Tar) but the repellent can only bind one of the receptor types (as the repellent amino acids often only can to Tsr) is equivalent to this model under *n*_*A*_ = *n*_Tsr_ + *n*_Tar_ and *n*_R_ = *n*_Tsr_ provided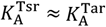. Specifically, for multiple ligands {L} binding multiple receptors {E}, the response function would be

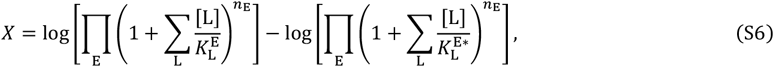

where 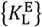 are the disassociation constants for ligand L binding receptor E in its inactive state and 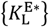 are the disassociation constants for the active state. If attractants tend to bind more types of receptor than do repellent or the attractant receptors are more abundant, this multi-chemoeffector multi-receptor variation will behave similarly to the simplified version used in this paper.

### Effect of fluid flows on nutrient plumes and microbe trajectories

**Fig. S1.**
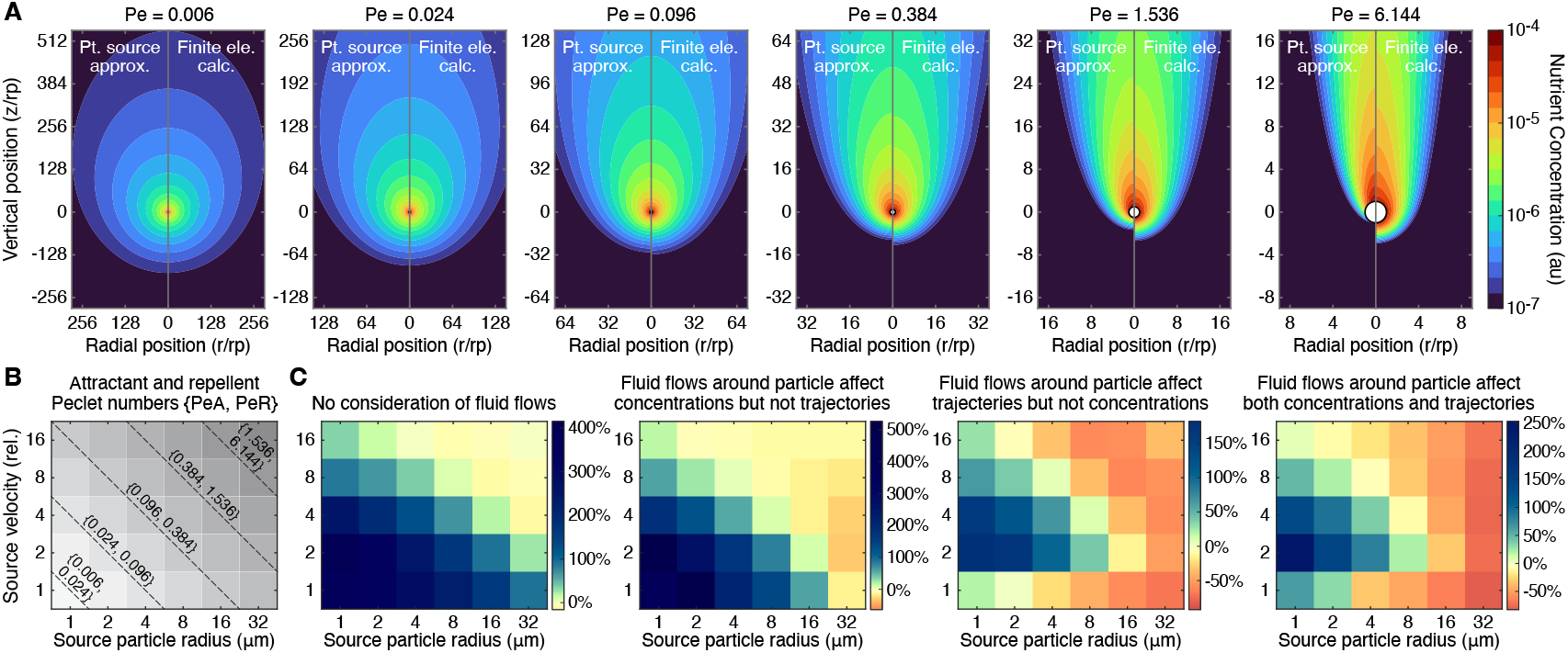
The differential strategy remains beneficial when accounting for the effects of fluid flows on concentration plumes and microbe trajectories. **A** Comparison of the point-source approximation used for calculating nutrients plumes throughout the Main Text (in which all nutrient is released from a point at the center of the source particle and there is no motion of the fluid around the source particle, Methods, left half of each plot) and results from numerically calculating nutrient plumes (using finite element analysis with a Dirichlet boundary condition at the surface of the particle and advection from fluid flows around the source particle incorporated into the calculation, Methods, right half of each plot) for a range of Péclet numbers (defined as pe_*i*_ = *r*_<_*v*_<_/*D*_*i*_). For each Péclet number, the same nutrient release rate is used for both the point source approximation and the finite element calculation. **B** Attractant and repellent Péclet numbers by source particle radius and relative velocity, assuming attractant and repellent diffusion coefficients of 1000 μm^2^/s and 250 μm^2^/s respectively and a microbe velocity of 6 μm/s. **C** Same as Fig. 3C (colormap showing the increase in intercept kernel for the differential vs purely attractant strategy) except, from left to right, (*i*) with no changes except the use of a coarse parameter grid, (*ii*) with concentration plumes calculated using finite element analysis instead of the point source approximation, (*iii*) with advection from fluid flows around the source particle accounted for when calculating microbe trajectories, and (*iv*) with both concentration plumes calculated using finite element analysis and the effect of fluid flows on microbe trajectories accounted for. Note that the same receptor binding affinities and cooperativities were used in calculating all four sets of results, but it is possible that the optimal values change when fluid flows are incorporated. These results should therefore be interpreted as lower bounds on the performance of the differential strategy and not as results from an optimized strategy.

#### Stochastic run-and-tumble simulations

**Fig. S2.**
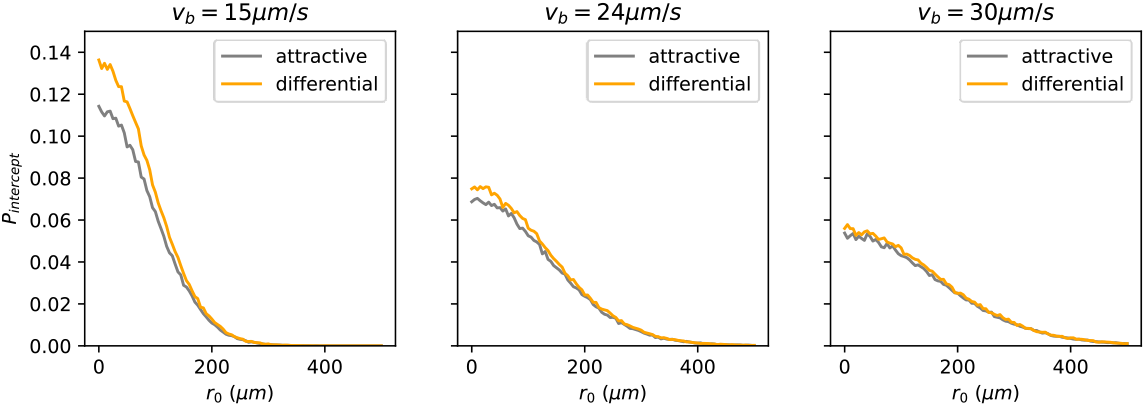
The differential strategy remains beneficial when accounting for the stochasticity of bacterial run-and-tumble chemotaxis. Probability to intercept the moving source, starting from distance *r*_0=_radially away and 500μm below the source (Methods). The plots show the probability based on 100,000 simulations from each *r*_0=_, spaced every 5μm up to *r*_=_ = 500μm. With intercept kernel 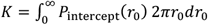, we find the advantage of differential strategy 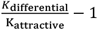 to be 16.0%, 7.36%, and 4.56% with increasing bacterial run speed *v*_*b*_ = 15μm/s, 24μm/s, and 30μm/s and correspondingly decreased sensitivity of the chemotactic response (Methods).

#### Marine microbe run-and-reverse simulations

**Fig. S3.**
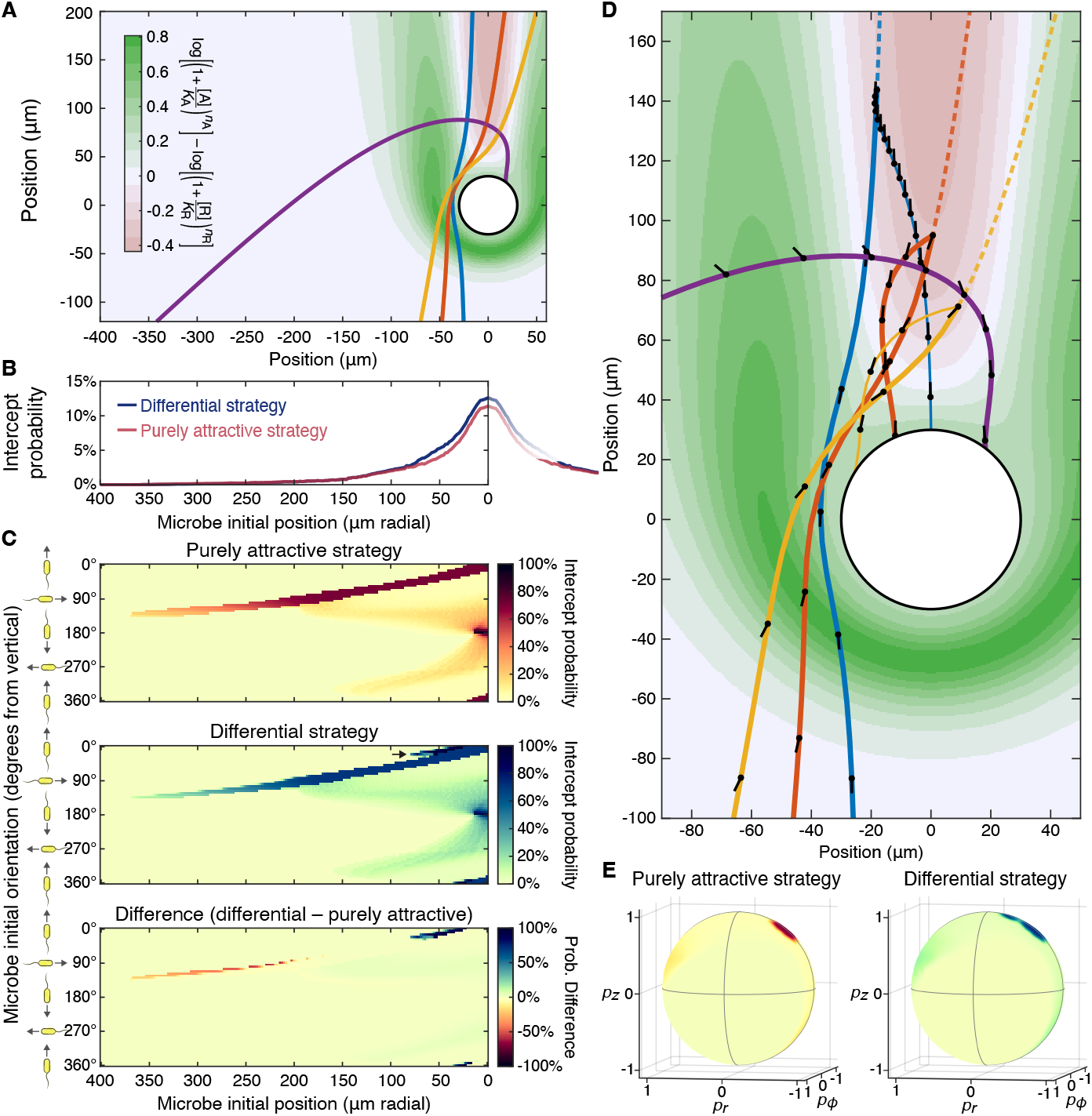
Run-and-reverse marine microbes also benefit from employing a differential strategy. **A** Particle (white circle) of radius 30 μm sinking at 200 μm/s and the differential response function (red-blue-green colormap) for an attractant and repellent each being released at 1 fmol/s^56^ (with *D*_*A*_ = 1000 μm^2^/s, *D*_R_ = 250 μm^2^/s, *n*_*A*_ = 4, *n*_R_ = 2, and *K*_*A*_ = *K*_R_ = 100 nM as in the Main Text). Due to the high Péclet numbers (pe_*A*_ = 6 and pe_*A*_ = 24), nutrient concentrations used to determine the response function are calculated using finite element analysis (i.e. without the point source assumption) as in Fig. S3 (Methods). Plotted on top of the response-function colormap are fluid streamlines around the particle in the particle frame (thin blue lines) and four microbe trajectories in the particle frame assuming the microbes swim at 150 μm/s and do not reverse (thick lines). Note that the microbes swim only in straight lines with the curvature of their trajectories coming from the fluid flows around the particle. **B** Probability of a microbe intercepting the source under the purely attractive (red) and differential strategies (blue) as a function of initial radial position (with a fixed initial vertical position of -100 μm) averaged over all possible initial orientations in 3D space weighted by the orientation-dependent vertical flux of microbes (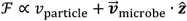, Methods). **C** For microbes starting with orientations in the x-z plane, the probability of successfully intercepting the particle under the purely attractive strategy (top), the probability of intercepting the particle under the differential strategy (middle), and the difference between those two probabilities (bottom) for run-and-reverse microbes with reverse probabilities dependent on the differential response function as described in the Main Text Methods. **D** Zoomed in version of **A** with trajectories drawn as thick lines until the point at which the “differential” microbes are most likely to reverse and then as thin lines for the trajectories for microbes that reverse then and microbes that do not reverse. The microbes’ positions at 200 ms increments are shown as a black dot with a black line extending opposite the direction in which it is swimming. Changes in the microbes’ orientations (except the reverse) are due to the fluid sheer and vorticity^39^, not active chemotaxis. **E** Probability of intercepting the source for all possible orientations in 3D space for a microbe starting at {*x*_0_, *y*_0_, *z*_0_} = {−*r*_0_, 0, *z*_0_} = {−60 μm, 0, −100 μm} using same colormaps as in **C**.

### Improvement in intercept kernel across particle velocity, density, and repellent release rate

**Fig. S4.**
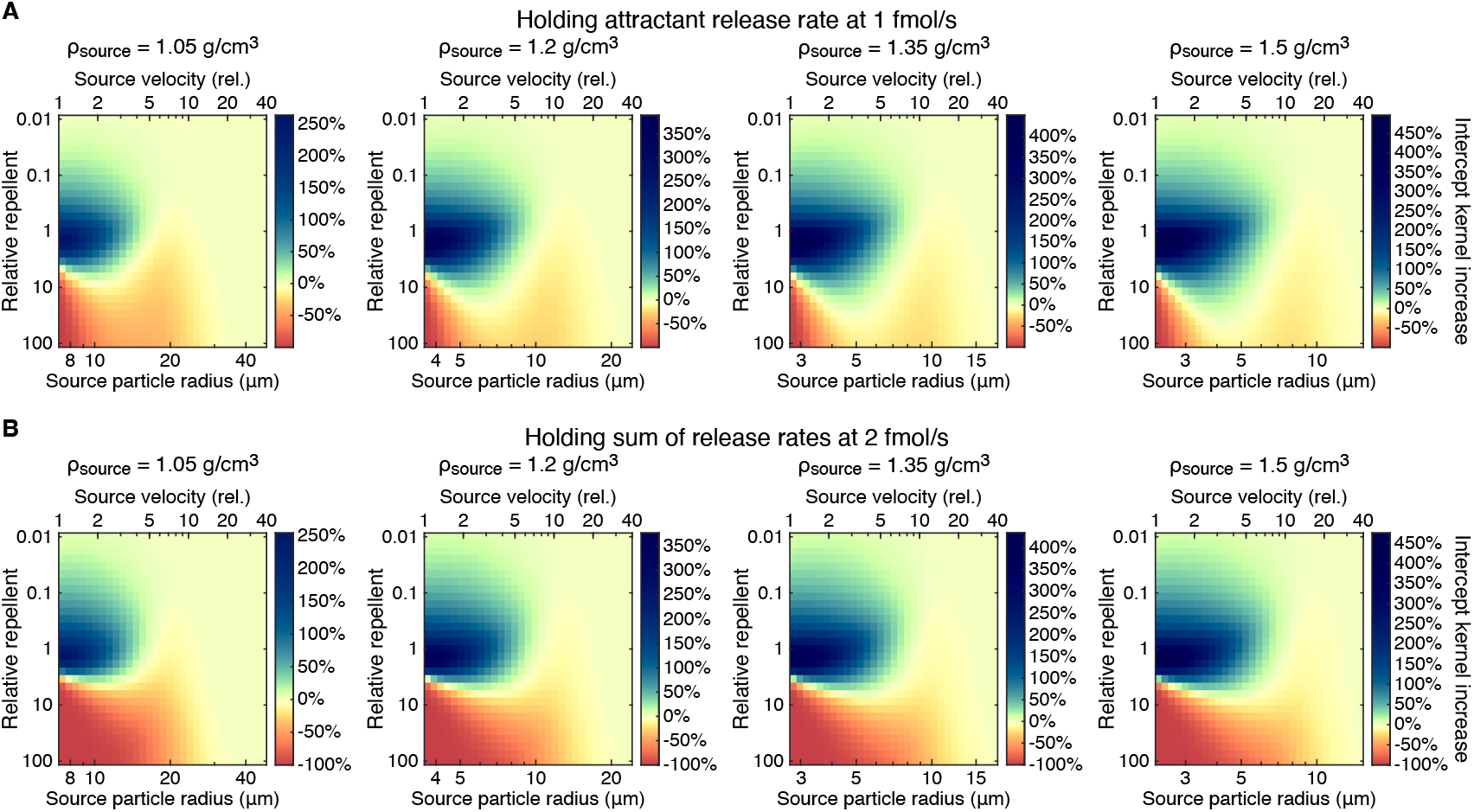
Improvement in the intercept kernel across source particle characteristics and relative repellent release rate. **A** Improvement in the intercept kernel as a function of source particle velocity (upper horizontal axis) and repellent release rate (relative to the attractant release rate of 1 fmol/s, vertical axis) for four source particle densities used to calculate the particle radii (lower horizontal axis) that correspond to the velocities (Methods). Across the full (three-dimensional) region shown, the mean intercept kernel under the differential strategy is 27% larger than the mean kernel under the purely attractive strategy. And across the subregion with relative repellent release rates between 0.1x and 10x, the mean kernel under the differential strategy is 78% larger. **B** Same as **A** except the attractant release rate is now also varied to maintain a total chemoeffector release rate of *s*_*A*_ + *s*_R_ = 2 fmol / s. Across the subregion with relative repellent release rates between 0.1x and 10x, the mean kernel under the differential strategy is 65% larger than under the purely attractive strategy.

